# Miniaturized and accessible functional ultrasound imaging system for freely moving mice

**DOI:** 10.1101/2025.04.24.650351

**Authors:** Jin Yang, Erlei Zhou, Jiejun Zhu, Tingting Zhu, Jianjian Zhao, Xiaoli Lin, Zheng Cao, Zihao Chen, Zhiwu An, Lei Sun, Wentao Wu, Zhihai Qiu

**Author notes:** These authors contributed equally to this work (Jin Yang, Erlei Zhou, Jiejun Zhu).

## Abstract

Functional ultrasound (fUS) imaging provides brain-wide activity maps with high spatiotemporal resolution and deep penetration, positioning it as a key technology for future non-invasive brain-computer interfaces (BCIs). Realizing this potential, particularly for chronic BCI applications requiring long-term monitoring in naturalistic settings, critically depends on significant system miniaturization to overcome the cost and complexity limitations of current platforms. Addressing this challenge, we present mini-fUS, a miniaturized, cost-effective fUS platform engineered for accessibility without compromising core performance for demanding neuroscience research. The system features a compact transmit-receiver, low-noise power supply, and high-speed data transfer, achieving pulse repetition frequencies up to 5 kHz with negligible jitter. Real-time GPU-accelerated beamforming and singular value decomposition (SVD) enable whole-brain activity mapping, demonstrated here in freely moving mice at up to 3.57 Hz with ∼100 µm spatial resolution and 15 mm penetration depth. Validated through recordings of brain activity during sensory stimulation, anesthesia, and behavior, this design defines a practical hardware-software framework for fUS. By significantly improving accessibility and enabling robust monitoring in mobile subjects, this work advances the development of fUS for both fundamental research and future BCI technologies, while clarifying essential fUS operational principles.

## Introduction

Understanding how brain-wide neural circuits orchestrate complex behaviors requires imaging tools capable of mapping activity across both cortical and subcortical regions with high spatiotemporal resolution, particularly in freely moving animals^[1, 2]^. Current techniques, however, present significant compromises: optical methods like multi-fiber photometry and miniaturized microscopy offer cellular or local circuit resolution but are limited in depth or field-of-view^[3, 4]^; electroencephalography (EEG)^[5]^ captures large-scale dynamics but lacks spatial precision; functional magnetic resonance imaging (fMRI)^[6]^ provides whole-brain coverage but is constrained by low temporal resolution and incompatibility with free movement; and while wide-field calcium imaging^[7]^ achieves high spatiotemporal resolution in mobile subjects, its shallow penetration depth restricts analysis primarily to superficial cortical layers. This leaves a critical gap in our ability to comprehensively monitor deep brain structures and inter-regional communication during naturalistic behaviors.

Functional ultrasound (fUS) imaging^[8]^ emerged as a powerful technology poised to bridge this gap^[9]^. By leveraging ultrafast plane-wave imaging^[10, 11]^ and sensitive Power Doppler measurements using Singular Value Decomposition (SVD) clutter filter^[12]^, fUS uniquely combines mesoscopic spatial resolution (∼100 µm), high temporal resolution (up to 10 Hz frame rates or higher), deep penetration (several centimeters), and a wide field-of-view, enabling non-invasive tracking of neurovascular dynamics across large brain volumes. Indeed, fUS has already provided valuable insights into brain function during sensory processing^[13-16]^, sleep^[17]^, cognitive tasks^[18-20]^, and disease states in diverse species^[21]^, from rodents to non-human primates and humans^[22-24]^. Furthermore, these capabilities make fUS a compelling technology for future non-invasive brain-computer interfaces (BCIs) that require detailed, brain-wide activity monitoring. However, the transformative potential of fUS for widespread neuroscience research, especially for applications demanding chronic monitoring in naturalistic settings like next-generation BCIs, has been significantly hampered by the substantial cost, complexity, and physical size of existing systems. Miniaturization is thus a critical bottleneck for unlocking these future applications. Typically built upon high-end clinical or research ultrasound platforms, these instruments often incorporate features unnecessary for preclinical neuroimaging, making them inaccessible to many laboratories and hindering the technique’s democratization.

To overcome this critical barrier and unlock the full potential of fUS for the neuroscience community, we developed and validated a miniaturized, integrated, and cost-effective 64-channel functional ultrasound (mini-fUS) system specifically engineered for high-fidelity brain imaging in freely moving mice. Our core innovation lies in a purpose-built design that prioritizes the essential requirements for neurovascular imaging—high sensitivity, sufficient spatiotemporal resolution, and robustness for mobile recordings—while eliminating costly and bulky components superfluous to this application. By leveraging compact semiconductor technology for transmit/receive functions (64 channels, 12-bit resolution, 50 MHz sampling, 2.5 ns delay precision for accurate beam steering) and integrating power supply and data transfer modules, we achieved a radical reduction in system footprint and cost compared to conventional platforms.

Crucially, miniaturization was achieved without sacrificing essential performance through systematic optimization of the data acquisition and processing pipeline. We implemented a highly efficient real-time beamforming strategy on a graphics processing unit (GPU) capable of handling compounded plane-wave imaging (e.g., 7 angles from -6° to 6°) at frame rates exceeding 1 kHz. Furthermore, we developed a fast singular value decomposition (SVD) clutter filter, employing optimized matrix operations that triple the processing speed compared to standard implementations. This integrated hardware-software co-design enables robust whole-brain functional imaging in freely moving mice at behaviorally relevant frame rates (e.g., 3.57 Hz using 200 compounded frames per Power Doppler image) with 100 µm resolution and up to 10 mm penetration. We complemented the hardware with user-friendly software, simplifying experimental workflows for researchers without specialized ultrasound expertise.

We demonstrate the system’s capabilities through comprehensive validation experiments. Whisker stimulation in head-fixed mice confirms its sensitivity to task-evoked hemodynamic responses. The isoflurane anesthesia experiment demonstrates the potential in pharmacological monitoring. Furthermore, we successfully recorded brain-wide activity patterns in freely moving mice during a pole test experiment, showcasing the system’s utility for studying brain states and behavior-neurovascular relationships under naturalistic conditions. By drastically lowering the technical and financial barriers to entry, our mini-fUS system represents a significant methodological advance, poised to democratize functional ultrasound imaging and empower broader investigation into the neural underpinnings of behavior across distributed brain circuits in health and disease.

## Results

### Comprehensive optimization design for mini-fUS

We developed a miniaturized functional ultrasound (mini-fUS) system optimized for accessible, high-performance brain imaging using low-cost integrated circuits (Fig. 1a). The modular architecture features two front-end (FE) boards, each controlling 32 Tx/Rx channels with 2.5-ns transmit delay precision via FPGA-based tri-state transmitters, enabling ultrafast plane-wave imaging (Fig. 1b, d). Echo signals are digitized at 50 M Samples/s with 12-bit resolution and transferred at 25 Gb/s using 8 lanes, each with a speed of 3.12 Gbps, supporting GPU-accelerated real-time beamforming and SVD filtering (Fig. 1e). The system imaged a 0.02-mm tungsten wire with a 0.17-mm lateral full width at half maximum (FWHM), demonstrating high spatial resolution and signal-to-noise ratio (Fig. 1f–h). A low-jitter (<500 fs RMS) 50-MHz clock ensures synchronized 64-channel acquisition at up to 5 kHz pulse repetition frequency. These results validate mini-fUS’s capability for precise, scalable fUS imaging, suitable for widespread neuroscience applications.

**Figure 1.**
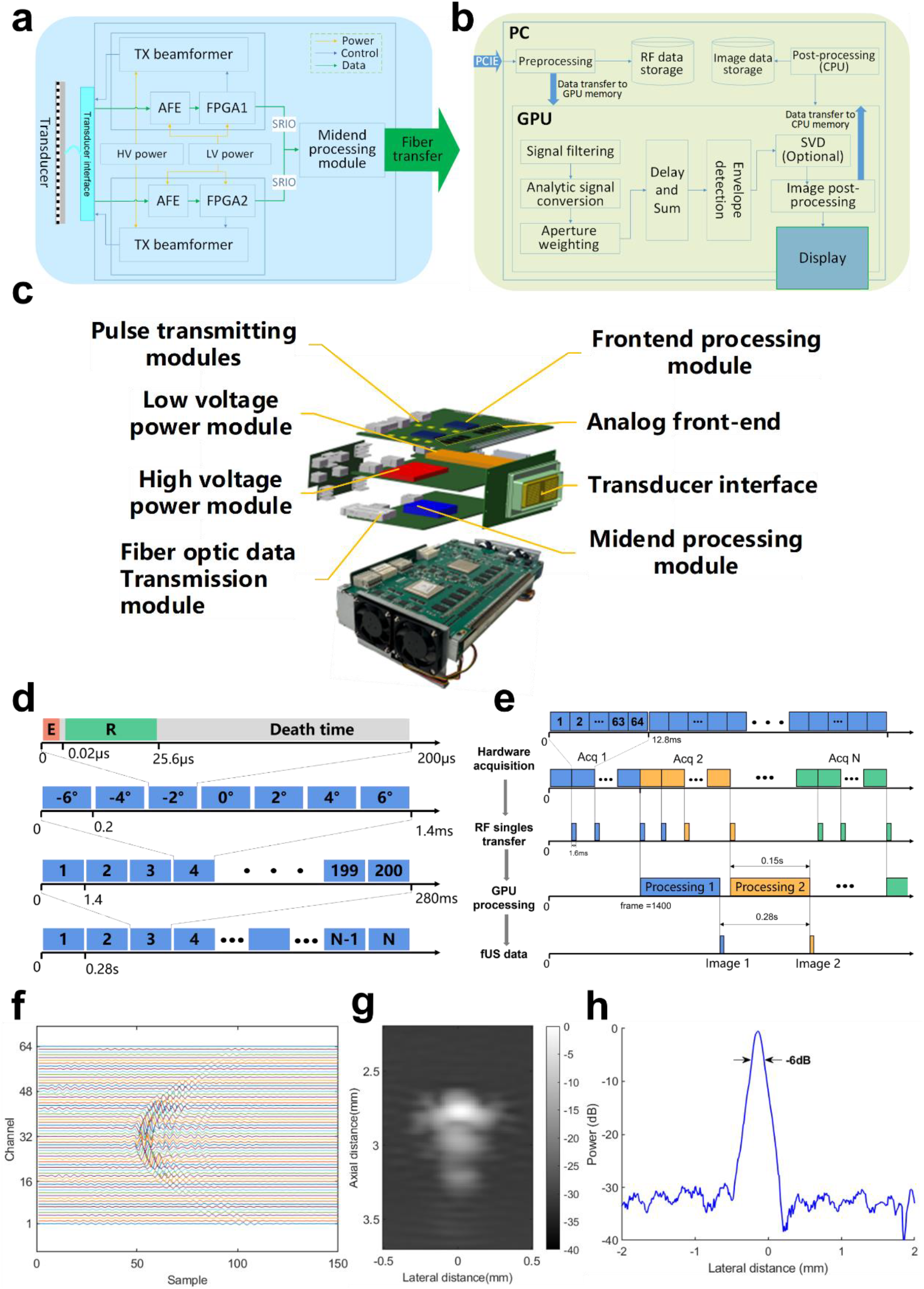
Miniaturized functional ultrasound imaging platform (mini-fUS) (a) The diagram of the ultrasound platform; (b) Real-time image reconstruction procedure; (c) 3D rendering of ultrasound imaging platform. Top: transmit and receive board (Tx & Rx). Middle: power supply board. Lower: post-processing board; (d) Ultrasound sequence to reconstruct a Power Doppler image; (e) fUS imaging acquisition and real time processing. (f) RF data of tungsten wire; (g) Ultrasound image of tungsten wire (d=0.02mm); (h) Lateral variation of tungsten wire image (FWHM=0.17mm)

### System function verification

We validated the imaging performance of the mini-fUS system using both controlled flow phantoms and in vivo mouse brain experiments.

First, to assess sensitivity to microvascular flow, we imaged a cornstarch solution flowing through polyethylene tubes of varying diameters driven by a cross-flow pump (Fig. 2a). Using a 15 MHz linear probe connected to the mini-fUS system, we successfully resolved flow within tubes as small as 100 µm in diameter (Fig. 2b, c), confirming the system’s suitability for microvascular imaging. We also observed that the Power Doppler (PD) signal intensity scaled directly with the programmed flow rate (Fig. 2d, e), demonstrating that the system provides quantitative information reflecting relative changes in blood flow, which is fundamental for functional neuroimaging studies inferring activity from hemodynamic changes.

**Figure. 2.**
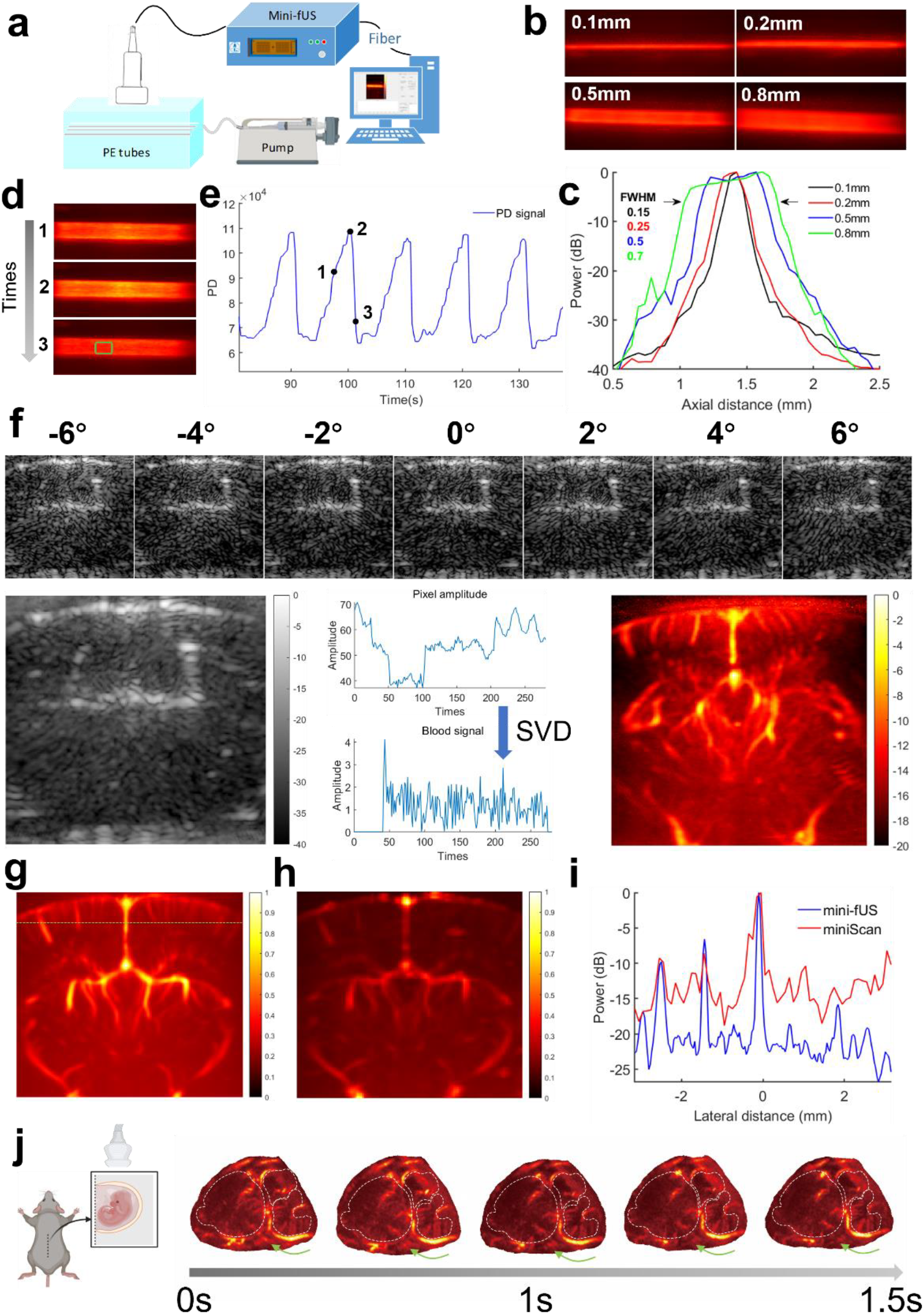
Performance validation of the mini-fUS system using phantoms and in vivo imaging. (a) Schematic diagram of the flow phantom setup designed to mimic microcirculation using tubes of varying diameters. (b) Representative PD images acquired by the mini-fUS system, showing resolved flow in tubes with inner diameters ranging from 0.1 mm to 0.8 mm. (c) Axial intensity profiles measured from the center of the tubes shown in (b). The Full Width at Half Maximum (FWHM) values were determined to be 0.15 mm, 0.25 mm, 0.5 mm, and 0.7 mm for the 0.1 mm, 0.2 mm, 0.5 mm, and 0.8 mm diameter tubes, respectively, indicating the system’s resolution capability. (d) Example PD images demonstrating the system’s ability to detect changes associated with periodically varying flow rates within the phantom. (e) Representative temporal trace of the PD signal intensity extracted from the region indicated by the green box in a flow image (e.g., from panel d). Higher signal intensity corresponds to increased flow rate. (f) Diagram illustrating the functional ultrasound (fUS) imaging sequence and processing pipeline implemented in the mini-fUS system. This includes multi-angle (−6° to +6°, 7 angles) plane wave emission, B-mode image compounding (example shown: 6.2 mm x 6.2 mm field of view, 40 dB dynamic range) for enhanced signal-to-noise ratio within 1.4 ms, and SVD filtering of the temporal signal *s*(*t*) from each pixel to isolate the blood flow components *s*_*b*_(*t*). The intensity of *s*_*b*_(*t*) is integrated over time (typically using 200 compounded frames acquired in 280 ms) to generate a PD image representing cerebral blood volume (CBV). (g) Representative coronal PD (CBV) map of a mouse brain acquired using the mini-fUS system. (h) Corresponding coronal PD (CBV) map from the same mouse brain acquired using the miniScan for comparison. (i) Lateral intensity profiles comparing vascular resolution across the cortical region indicated by the green dashed line in (g) and (h). (j) Representative coronal PD (CBV) map of a mouse embryo brain at embryonic day 18.5 (E18.5), acquired with the mini-fUS system. Green arrows highlight resolved major vascular structures in the forebrain, midbrain, and hindbrain.

Next, we applied the mini-fUS system for in vivo brain imaging in anesthetized mice. We employed a sequence typical for functional imaging: transmitting plane waves at seven distinct angles (−6° to +6°), compounding the resulting B-mode images, and accumulating 200 such compounded frames to generate a single high-sensitivity PD image representing cerebral blood volume (CBV) (Fig. 2f). This process effectively isolates the Doppler signal *s*_*b*_(*t*) arising from moving red blood cells from tissue echo signal *s*(*t*) including larger, lower-frequency tissue motion signal using SVD based clutter filtering, integrated into our real-time processing pipeline.

To benchmark the performance against established platforms, we compared PD images of the same mouse brain acquired sequentially with our mini-fUS and miniScan (a fUS platform based on Vantage system)^[25]^ (Fig. 2g, h). Qualitative comparison revealed detailed visualization of cortical and subcortical vasculature with both systems. Quantitative analysis of a lateral intensity profile across cortical vessels (Fig. 2i, corresponding to green dashed line in Fig. 2g) indicated that the mini-fUS system achieved comparable or potentially improved spatial resolution, characterized by narrower peak widths corresponding to individual vessels and potentially lower side-lobe artifacts, enabling clear differentiation of adjacent micro vessels.

To further highlight the system’s versatility, we imaged the complex vasculature within a developing mouse embryo brain (Fig. 2j). The mini-fUS system clearly resolved the primary vascular networks spanning the forebrain, midbrain, and hindbrain (Fig. 2j, green arrows), demonstrating its utility for detailed anatomical visualization beyond the adult brain. Taken together, these phantom and in vivo results validate that our optimized, miniaturized mini-fUS system achieves high-fidelity vascular imaging performance on par with conventional, more complex ultrasound systems, confirming its suitability for demanding research applications.

### Functional experiments verified by Task-evoked brain activation experiment Pharmacological experiment

To evaluate the capability of the mini-fUS system for detecting transient, localized brain activity, we imaged cerebral hemodynamics during sensory stimulation in awake, head-fixed mice. We performed fUS acquisitions at a frame rate of 3.57 Hz while applying periodic stimulation to the mouse’s left whiskers (experimental timeline depicted in Fig. 3a).

**Figure 3.**
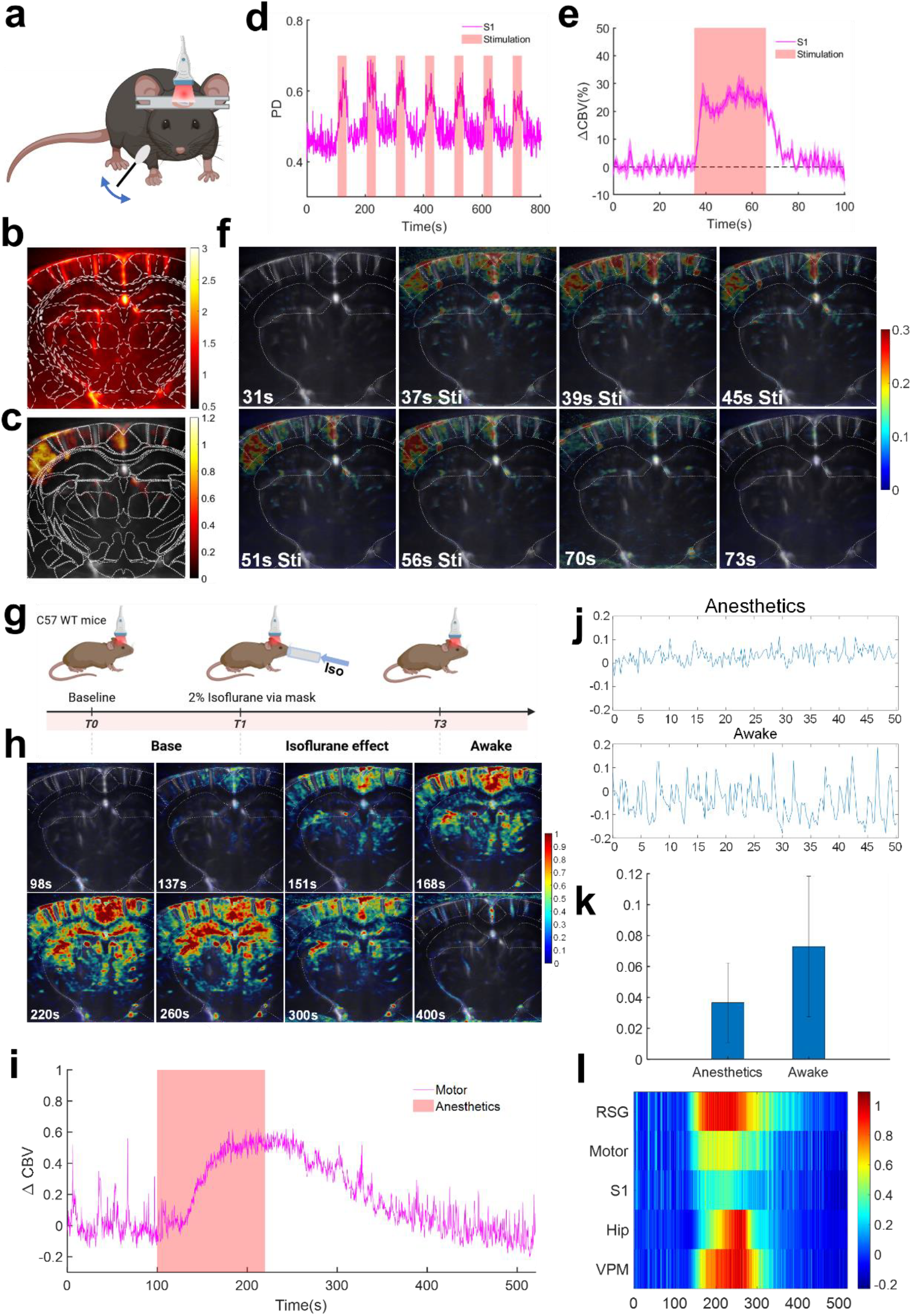
Applications of functional ultrasound (fUS) imaging with mini-fUS. (a) Schematic of the whisker stimulation experiment in awake, head-fixed mice. (b) Power Doppler image of the mouse brain (coronal plane, stereotaxic coordinates β = −2.06 mm). (c) Activation map from right whisker stimulation, showing voxel-wise correlation coefficients between Power Doppler signals and the stimulus pattern, with activation localized to the contralateral primary somatosensory barrel cortex (S1bf). (d) Power Doppler signal in S1bf during whisker stimulation (35 s baseline, 30 s on [red shade], 35 s off, repeated 7 times). (e) Average change in cerebral blood volume (ΔCBV) in S1bf, showing a ∼20% increase in Power Doppler signal during stimulation. (f) Spatiotemporal whole-brain activity during stimulation, with Power Doppler signal changes calculated per pixel every 3 s. (g) Schematic of the isoflurane anesthesia experiment (2% dose). (h) Brain activity during anesthesia, showing increased cerebral blood volume across cortex, hippocampus, and deeper regions. (i) ΔCBV response in the motor cortex during anesthesia. (j) Comparison of ΔCBV in the motor cortex between awake and anesthetized states. (k) Average oscillation amplitude of CBV signals in the motor cortex, showing low-frequency oscillations with reduced amplitude during anesthesia (3.7 ± 2.6%) compared to the awake state (6.4 ± 4.4%).

Analysis across seven stimulation trials revealed a robust and localized hemodynamic response. We generated an activation map by calculating the voxel-wise correlation coefficient ***r*** between the Power Doppler signal time course and the temporal pattern of the whisker stimulation blocks. This map showed significant activation predominantly localized to the contralateral primary somatosensory barrel cortex (S1bf), consistent with the known functional neuroanatomy (Fig. 3c).

Critically, the mini-fUS system demonstrated high sensitivity, reliably detecting hemodynamic increases within S1bf even on single trials (representative single-trial response shown in Fig. 3d). Averaging the response across trials revealed a steady increase in blood volume during the stimulation period, with the Power Doppler signal increasing by approximately 20% from baseline (Fig. 3e). Furthermore, the combination of high temporal resolution and wide field-of-view allowed us to visualize the spatiotemporal evolution of the activity pattern across the imaged brain slice, capturing the transition from the resting state to the stimulus-evoked activated state (Fig. 3f). These results demonstrate the mini-fUS system’s effectiveness in non-invasively monitoring task-related neural activity via neurovascular coupling in awake, behaving animals.

### Mini-fUS Monitors Hemodynamic Effects of Isoflurane Anesthesia

We also investigated the effects of 2% isoflurane anesthesia on cerebral hemodynamics (Fig. 3g). Following a 100 s awake baseline recording, mice underwent 120 s of anesthesia and a subsequent 300 s recovery period. Whole-brain Power Doppler analysis revealed that isoflurane administration initially increased cerebral blood volume (CBV), observed as an increase in Power Doppler signal intensity that began in the cortex and spread progressively to deeper regions. This elevated CBV normalized gradually upon cessation of anesthesia as the mice awoke, consistent with isoflurane’s known vasodilatory effects.

While mean CBV increased, the temporal dynamics of the Power Doppler signal were significantly altered. Specifically, cortical signal fluctuations were visibly reduced during stable anesthesia compared to the awake state (representative traces, Fig. 3i). Quantitative analysis comparing stable signal periods confirmed this: the amplitude of fluctuations of the cortical Power Doppler signal was significantly lower during anesthesia (3.7 ± 2.6%) compared to the awake state (6.6 ± 4.4%) (Fig. 3j, k). This likely reflects the suppression of spontaneous neural activity under anesthesia.

These findings underscore the mini-fUS system’s capability to track both drug-induced changes in overall blood volume and concurrent alterations in hemodynamic variability, demonstrating its versatility for neuropharmacological investigations alongside task-evoked activity studies.

### Mini-fUS Captures Behavior-Correlated Brain Activity During Freely Moving Pole Test

To investigate the relationship between behavior and brain activity in an unconstrained setting, we utilized the mini-fUS system to monitor CBV dynamics in freely moving mice performing a pole climbing task. A custom-designed lightweight (approx. 2g) headstage incorporating the 64-element micro-probe was affixed to the mouse’s head, enabling fUS imaging during unrestricted movement (Fig. 4a). In the task, mice placed at the top of a vertical pole were allowed to climb down autonomously; fUS data were recorded across five repeated trials.

**Figure 4.**
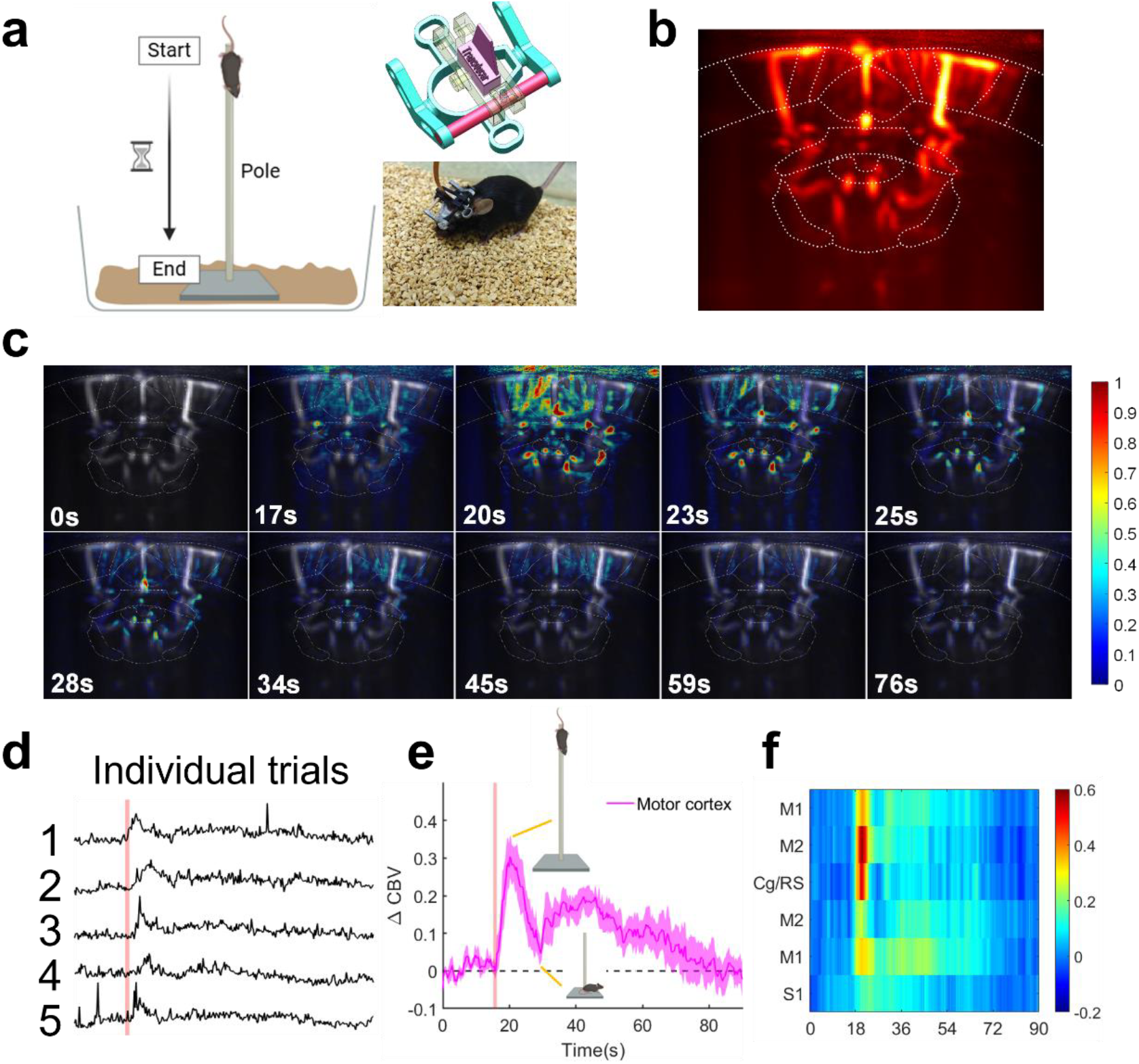
Freely moving application of mini-fUS in the pole test. (a) Schematic of the pole test experiment. Top right: Lightweight (2g) headstage designed for freely moving mice, with a probe holder adjustable along the β-axis for imaging position selection. Bottom right: Photograph of a mouse with the headstage during fUS imaging. (b) fUS image of the mouse brain in a freely moving state. (c) CBV signal across the whole brain during the pole test, showing significant activation in the motor cortex. (d) CBV signal in the motor cortex during the experiment, which involved free crawling, lifting mice to the pole’s top, and autonomous climbing to the ground (repeated 5 times). (e) Average CBV signal in the motor cortex, peaking when mice were at the pole’s top, gradually decreasing as they descended, and slightly rising before returning to baseline. (f) CBV signals in six brain regions during the pole test, corresponding to regions in (b) from left to right: primary motor cortex (M1), secondary motor cortex (M2), cingulate/retrosplenial cortex (Cg/RS), M2, M1, and primary somatosensory cortex (S1).

We observed significant task-related activation, particularly in motor control areas. Analysis revealed robust increases in CBV predominantly within the primary and secondary motor cortices (M1/M2) during active climbing (representative activation map, Fig. 4c). The temporal dynamics of the CBV signal closely tracked the behavioral sequence within each trial as illustrated by representative traces comparing CBV and movement timing in Fig. 4d. Averaging the CBV signal from the motor cortex across the five trials (Fig. 4e) showed peak activation near the onset of descent from the pole top, followed by a gradual decline as the mouse descended. A transient signal modulation occurred after the mouse reached the ground, before returning to baseline levels.

Furthermore, analysis across multiple cortical regions revealed distinct, region-specific activity patterns associated with the task. Figure 4f displays the average time courses of CBV signals from representative areas, including bilateral motor (M1, M2), cingulate/retrosplenial (Cg/RS), and somatosensory (S1) cortices (assuming this reflects the labels), highlighting differential engagement during the pole climb.

These results successfully demonstrate the capability of the mini-fUS system, facilitated by its miniaturized components and lightweight headstage design, to capture dynamic, behavior-locked brain activity across distributed cortical regions simultaneously in freely moving animals.

## Discussion

Functional ultrasound (fUS) imaging offers a powerful window into brain activity by monitoring neurovascular coupling with exceptional spatiotemporal resolution. However, its widespread adoption in neuroscience has been significantly hindered by the cost, size, and complexity of conventional systems, which are often adapted from multi-purpose clinical platforms. To address this critical gap, we developed a miniaturized fUS (mini-fUS) system based on a design philosophy prioritizing balanced performance and accessibility. Our approach focused on optimizing the essential elements for high-fidelity neurovascular imaging—specifically ultrafast plane-wave scanning and efficient GPU-accelerated computation (Fig. 1a, b)—while leveraging cost-effective semiconductor technology. This streamlined design demonstrates that robust Power Doppler imaging is achievable even with a 12-bit acquisition system when coupled with optimized processing, simplifying fUS implementation. The system’s modular hardware, built around 32-channel Tx/Rx units scalable to 64 channels (and potentially beyond), provides flexibility for 2D imaging and establishes a foundation for future 3D applications. Furthermore, the accompanying GPU-based toolkit, providing real-time beamforming and SVD clutter filtering, is envisioned as a potential open-source contribution to standardize and streamline fUS data processing across different platforms.

The mini-fUS system’s performance was rigorously validated. We demonstrated high spatial resolution, capable of resolving cortical microvessels with an FWHM of approximately 102 μm (Fig. 2g, i). Crucially, the system exhibited high sensitivity, sufficient to detect hemodynamic responses to whisker stimulation on a single-trial basis in awake mice, showing a ∼20% Power Doppler signal increase in the relevant somatosensory cortex (Fig. 3a–f). This sensitivity highlights its potential for capturing subtle or transient neural events, potentially offering advantages over fMRI in specific contexts demanding higher temporal resolution. We further demonstrated its utility in pharmacological studies, revealing both the expected isoflurane-induced increase in cerebral blood volume (CBV) across cortex and hippocampus, and a concurrent, significant reduction in the temporal fluctuations of the cortical Power Doppler signal from 6.6 ± 4.4% (awake) to 3.7 ± 2.6% (anesthetized) (Fig. 3g–k), likely reflecting suppressed neural activity. Importantly, equipped with a lightweight (2g) headstage and 64-element micro-probe, mini-fUS enabled artifact-free imaging during a demanding pole climbing task in freely moving mice, successfully capturing dynamic CBV peaks correlated with movement initiation in the motor cortex and distinct region-specific activity patterns (Fig. 4). These diverse applications collectively underscore the mini-fUS system’s versatility and robustness for task-evoked, pharmacological, and behavioral neuroscience research.

Compared to existing fUS platforms, mini-fUS represents a significant step towards democratization, offering substantial reductions in size and cost while maintaining competitive performance in terms of penetration depth, resolution, and potentially enhanced signal-to-noise ratio through optimized processing. Its proven compatibility with awake and freely moving states opens possibilities for chronic studies and applications in behaving animals, potentially including non-human primates where compact, robust systems are advantageous. It is important to note that while the current mini-fUS system, particularly the associated electronics and cabling, is not yet sufficiently miniaturized to be fully carried by smaller animals like mice or monkeys as a “backpack” during large-scale navigation, our work fundamentally demonstrates the feasibility and potential of this miniaturization pathway. The achieved reduction in size and complexity using integrated electronics establishes a crucial foundation for future iterations aimed at further significant miniaturization. Such future developments could ultimately enable continuous, untethered brain-wide monitoring during truly unrestricted exploration in larger environments. While the current implementation provides high-resolution 2D imaging, a primary limitation, the modular architecture is explicitly designed to facilitate expansion towards multi-probe configurations for volumetric (3D) imaging in future work. By substantially lowering the technical and financial barriers to entry, mini-fUS has the potential to transition fUS from a niche technique to a routinely accessible laboratory tool. This dedicated, optimized system can empower a broader range of research, including task monitoring, high-throughput drug screening, and the development of novel brain-machine interfaces, thereby fostering scalable, high-impact neuroscience discoveries.

## Conclusion

In summary, we have developed mini-fUS, a miniaturized, cost-effective, and high-performance functional ultrasound system specifically optimized for the demands of neuroscience research. By successfully balancing advanced imaging capabilities with accessibility and ease-of-use, this system overcomes significant barriers that have limited the broader adoption of conventional fUS technology. We have demonstrated its robustness and versatility across task-evoked activation, pharmacological monitoring, and challenging freely moving behavioral paradigms. Mini-fUS represents a significant step towards democratizing functional brain imaging, poised to empower a wider range of researchers and accelerate discoveries in neural circuit function, behavior, and neuropathology.

## Method

### Mice

All experiments were performed on C57BL/6J mice obtained from Bestest (Zhuhai, China). Mice were housed with a regular dark–light cycle and fed a standard diet and water *ad libitum*. All mouse procedures were performed following the guidelines of the Committee of Use of Laboratory Animals, Guangdong Institute of Intelligence Science and Technology, China.

### Surgery

Eight-week-old C57BL/6 mice were anesthetized with a combination of anesthetics (a mixture of Zoletil 50 and xylazine 10μL/g) and secured in a head stereotaxic apparatus. After shaving the head, the scalp was thoroughly disinfected with 75% ethanol and incised along the midline to expose the skull. The fascia covering the cranial surface was carefully removed to fully reveal the target area between the bregma and lambda. A handheld micro-drill (RWD 78001, China) was used to slowly drill into the predetermined area, with sterile saline continuously applied to the drilling site to prevent overheating and potential brain tissue damage. Once the boundary was completely penetrated, sterile saline was applied to the drilled area for approximately 3 minutes to soften the bone plate, after which the skull was carefully removed, ensuring the dura mater remained intact. A transparent polymethylpentene (TPX) membrane was placed over the craniotomy site to serve as an acoustic window for ultrasound imaging. The edges of the window were sealed with sterile medical adhesive and reinforced with dental cement (Super-Bond C&B, Japan). If repeated imaging or probe stabilization was required, a custom head-fixation mount was also implanted. Postoperatively, cefotaxime sodium (5 μg/g) was administered intraperitoneally for 3 consecutive days to prevent infection. The mice were allowed to recover undisturbed for 7 days before undergoing functional ultrasound imaging experiments.

### Functional ultrasound imaging

#### Data acquisition

Data was acquired using a linear ultrasound probe composed of 64 piezoelectric transducers with a center frequency of 15 MHz and a 100μm pitch. In the head-fixed experiments, the probe was mounted on a three-dimensional translation stage to achieve precise positioning and scanning. In the freely moving experiments, the probe was attached to a custom head-mounted holder on the mouse, allowing it to slide along a guide rail to position the brain slice accurately. fUS is performed by acquiring 200 compound images, each one from 7 plane wave scanning title from -6 º to 6 º at PRF of 5000 Hz. To enhance real-time computing capabilities and increase temporal resolution, RF data is processed on a professional-grade graphical computing card (Nvidia RTX 4090) was integrated, enabling a recording frequency of 3.57 Hz.

#### Beamforming

For a plane wave with a steering angle of α, the RF-data is beamformed using the Delay and Sum (DAS) algorithm, the beamforming equation is as follows:

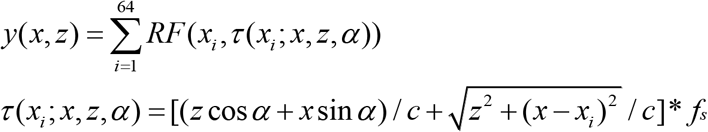

Here, *y*(*x,z*) represents the echo signal intensity at pixel (*x,z*), *τ* is the time index, *f*_*s*_ is the sampling frequency, and *c* is the speed of sound. By processing all pixels, a single plane wave imaging (PWI) ultrasound image can be obtained. This process will generate 7*200 PWIs in total.

After beamforming, the image SNR is enhanced using coherent plane wave compounding. This involves sequentially stacking plane wave images acquired at 7 different angles (ranging from -6º to 6º) to produce a single compounded plane wave image. Each functional ultrasound (fUS) sequence will generate 200 compounded plane wave images.

After beamforming, coherent plane wave compounding (CPWC) is used to enhance the image signal-to-noise ratio. Specifically, this involves sequentially summing PWIs acquired at 7 different angles (ranging from -6º to 6º) to create a single compounded plane wave image (CPWI). Each functional ultrasound (fUS) sequence will generate 200 CPWIs.

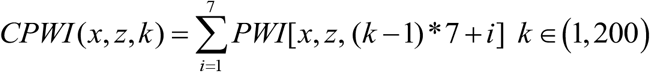

#### SVD

To perform the Power Doppler image, we use a singular value decomposition (SVD) filter to extract blood flow signals. Compared to traditional high-pass filters, the SVD filter is more sensitive to micro blood flow.

Considering the matrix 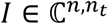 where the beamformed and compounded frames have been stacked in columns, the SVD of *I* is:

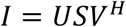

Where 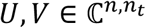 are unitary matrices containing the spatial and temporal singular vectors in there columns respectively, 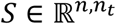 represents a diagonal matrix containing _*t*_ singular values *s*_*i*_ ∈ ℝ^+^ of *I* in its diagonal arranging in descending order 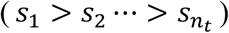, *n* is the number of pixels and *n*_*t*_ is the number of compound images.

The SVD subspaces are linked to the three main components of ultrasound data: tissue, blood, and noise signals. The tissue signal, which is highly coherent both in space and time, occupies the low-rank subspaces with high singular values. The uncorrelated noise occupies the subspaces with low singular values while blood components lie between the two subspaces. Therefore, we can retain blood signal *s*_*b*_(*t*) based on the magnitude of the singular values while removing tissue and noise signals. And the Power Doppler image can be performed as:

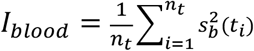

### GPU acceleration

To perform fUS imaging, the processes of beamforming and SVD filtering are implemented on the GPU platform. For beamforming, parallel processing over multiple GPU cores required a precise assignation of all tasks in a grid composed of blocks. Each block of this grid is assigned to a different thread computed by a specific core. For SVD processing, a fully parallel CUDA matrix operation library is used to accelerate SVD filtering. Power Doppler images can be computed in real-time (3.57 frame per second). This process function should run in a computer equipped with a NVIDIA GPU. And CUDA runtime software must be also installed.

### Experiment

#### Whisker stimulation experiment

In this experiment, the mouse was placed in a mouse holder on a thermostatically controlled heating pad set to 37°C, with its head securely fixed. Coupling gel was applied to the cranial window. The probe was adjusted to the imaging position (β -2.06 mm, 3 mm from the brain surface) with an imaging field of view of 6 mm by 6 mm. The probe position was further adjusted to ensure that the imaging field covered the left hemisphere, specifically the left primary somatosensory cortex. During the fUS acquisition, a cotton swab was used to continuously stimulate the mouse’s right whiskers, synchronized with the sampling times of the ultrasound scanner. The stimulation protocol consisted of 35 seconds of baseline, followed by 30 seconds of stimulation and 35 seconds of rest, repeated seven times.

### Isoflurane-Anesthetized experiment

In this experiment, anesthesia was induced using 2% isoflurane administered via a mask. The mice were secured in a mouse holder with its head fixed, and erythromycin ointment was applied to protect their eyes. A thermostatically controlled heating pad was set to maintain body temperature at 37°C. An air-free ultrasound coupling gel was applied to the cranial window, and the probe was adjusted to the imaging position (β - 2.06 mm, 3 mm from the brain surface). The experimental protocol was as follows: 100 seconds of baseline data were collected, followed by 120 seconds of recording during anesthesia induction after turning on the vaporizer, and then 300 seconds of recording during the recovery process after turning off the vaporizer. This procedure was repeated for three sessions.

### Freely moving pole test experiment

In this experiment, air-free ultrasound coupling gel was filled into the central container of the head-fix until the container was completely filled. The design of the central container aims to prevent the loss of coupling gel during the mouse’s movement. Similarly, coupling gel was injected into the probe holder to avoid any residual air after installing the probe. The miniature ultrasound probe was then mounted into the probe holder. The mini-fUS was activated to verify the quality of the blood flow images, and the probe holder was slid along the guide rail to select the scanning position. Once the scanning position was confirmed, the screws on the probe holder were tightened to secure it in place.

During the preparation phase of the experiment, a straight pole was placed in the mouse’s living cage, and the mini-fUS was activated to allow the mice to move freely with the probe for 30 minutes to acclimate to the experimental environment. The experimental protocol was as follows: 100 seconds of free movement signals were recorded as a baseline. The mouse was then lifted to the top of the pole and released to climb down freely. The time taken for the mouse to reach the top and return to the bottom was synchronized with the ultrasound scanner, and cerebral blood flow signals were continuously recorded after the mouse left the pole (total 100s). This procedure was repeated 10 times, and data sets with motion artifacts were excluded.

### Analysis

#### Registration

We first evaluate the collected fUS data and exclude any data affected by image artifacts. These artifacts may be caused by factors such as abnormal mouse movement, uneven coupling agent, or the presence of bubbles. To achieve accurate functional analysis, it is necessary to register the collected cerebral blood flow data with the Allen Brain Atlas. This process requires us to acquire a stable cerebral blood flow image at the beginning of the experiment and align it with a reference cerebral blood flow image that has already been registered to the standard brain atlas. This ensures that the cerebral blood flow data collected in subsequent experiments automatically aligns with the standard atlas.

#### General Analysis Methods

Based on the principle of neurovascular coupling, the fundamental analysis method of fUS involves assessing the changes in blood volume across different brain regions to determine their activity states. Since the PD signal intensity is proportional to blood volume, we can calculate the average PD signal intensity for each brain region and plot the trace of PD signal changes over time. To further enhance the analysis of brain activity, we calculate the rate of blood volume change (ΔCBV), expressed as the percentage change of the PD signal relative to a baseline value for each brain region. Additionally, we analyze the characteristics of blood volume changes on a whole-brain scale, which is a significant advantage of fUS. The specific method involves calculating the percentage change of the PD signal *I(x,z,t)* for each pixel relative to the baseline, denoted as *ΔI(x,z,t)*, and applying a temporal window (1-3s) to smooth *ΔI(x,z,t)*. This reduces the interference from discrete errors and highlights responsive regions. The resulting data allows for a more comprehensive analysis of whole-brain activity during experiments, facilitating the rapid identification of specific brain regions, functional connectivity, and other features.

## Acknowledgments

This work was supported in part by the National Key Research and Development Program of Ministry of Science and Technology of China (2023YFC2410900), National Natural Science Foundation of China (32371151), Hong Kong Research Grants Council Collaborative Research Fund (C5053-22 GF), General Research Fund (15126524 and 15224323), and internal funding from the Hong Kong Polytechnic University (G-SACD), Research Center for Non-invasive Brain Computer Interface (1-CE0M), Research Institute of Smart Ageing (1-CDJM), and Independent Deployment Project of Institute of Acoustics, Chinese Academy of Sciences (JCQY202413).

## Author Contributions

Conceptualization: ZHQ, JY, JJZ; Methodology: JY, ELZ, WTW, ZC, ZWA; Experiment: JY, TTZ, XLL, JJZ; Data analysis: JY, ZHC; Manuscript writing: ZHQ, JY; Funding acquisition: ZHQ, LS; Supervision: ZHQ, LS.

## Competing Interest Statement

The authors declare that they have no competing interests. Data and materials availability: All data are available in the main text or the supplementary materials.

## Notes

### Competing Interest Statement

The authors have declared no competing interest.

